# Insights into the salivary *N*-glycome of *Lutzomyia longipalpis*, vector of visceral leishmaniasis

**DOI:** 10.1101/2020.06.03.132746

**Authors:** Karina Mondragon-Shem, Katherine Wongtrakul-Kish, Radoslaw P. Kozak, Shi Yan, Iain Wilson, Katharina Paschinger, Matthew E. Rogers, Daniel I. R. Spencer, Alvaro Acosta-Serrano

## Abstract

During *Leishmania* transmission sand flies inoculate parasites and saliva into the skin of vertebrates. Saliva has anti-haemostatic and anti-inflammatory activities that evolved to facilitate bloodfeeding, but also modulate the host’s immune responses. Sand fly salivary proteins have been extensively studied, but the nature and biological roles of protein-linked glycans remain overlooked. Here, we characterised the profile of *N*-glycans from the salivary glycoproteins of *Lutzomyia longipalpis*, vector of visceral leishmaniasis in the Americas. *In silico* predictions suggest half of *Lu. longipalpis* salivary proteins may be *N*-glycosylated. SDS-PAGE coupled to LC-MS analysis of sand fly saliva, before and after enzymatic deglycosylation, revealed several candidate glycoproteins. To determine the diversity of *N*-glycan structures in sand fly saliva, enzymatically released sugars were fluorescently tagged and analysed by HPLC, combined with highly sensitive LC-MS/MS, MALDI-TOF-MS, and exoglycosidase treatments. We found that the *N*-glycan composition of *Lu. longipalpis* saliva mostly consists of oligomannose sugars, with Man_5_GlcNAc_2_ being the most abundant, and a few hybrid-type species. Interestingly, some glycans appear modified with a group of 144 Da, whose identity has yet to be confirmed. Our work presents the first detailed structural analysis of sand fly salivary glycans.

## Introduction

Sand flies are small insects that can transmit bacteria and viruses^1,2^, but are known mainly as vectors of leishmaniasis, a disease that threatens 350 million people worldwide^3^. When female sand flies feed, they inject a saliva comprised of molecules that facilitate the ingestion of blood, and modulate the host immune system and pathogen transmission^4,5,6^. These effects have led researchers to explore the potential of insect salivary molecules as markers of biting exposure^5,7^ (to determine risk of disease), or even as components of vaccines against leishmaniasis^8^. Of the many types of molecules that make up saliva, most research has focused on the proteins; here, we have investigated the glycans that modify these proteins.

In most eukaryotic cells, the addition of glycans to proteins is a highly conserved and diverse post-translational modification. The most common types of protein-linked glycans are *N*-linked (attached to asparagine residues in the sequon Asn-X-Thr/Ser), and *O*-linked (attached to serine or threonine residues). Glycoconjugates display a wide range of biological roles, from organism development to immune system functions against pathogens^9^. One study has addressed the types and roles of glycans in insects using the model fruit fly, *Drosophila melanogaster*. In this species, biological functions have been attributed to different glycan classes, such as morphology and locomotion (*N*-linked glycans), or cell interaction and signalling (*O*-linked glycans)^10^.

Glycans may have special relevance in the saliva of medically important arthropods, because of the fundamental role this biological fluid plays during pathogen transmission. For instance, African trypanosomes, tick-borne pathogens, arboviruses and malaria are all harboured in the salivary glands of their respective vectors, and are co-transmitted with saliva through the bite. In contrast, *Leishmania* parasites are transmitted by regurgitation from the fly’s midgut, where infectious stages reside, and contact with saliva occurs in the host at the bite site^11^. People living in leishmaniasis-endemic regions are constantly exposed to the saliva of uninfected sand flies, triggering immune responses that may later influence parasite infection^12^. The immunogenicity of salivary glycan structures and their interaction with specific immune cells could have different effects for each disease.

There are some reports describing the presence of salivary glycoproteins in sand flies through *in silico* and blotting analyses^13-19^; however, to our knowledge no detailed structural studies have been published to date. Therefore, we set out to identify the salivary glycoproteins in the sand fly vector species *Lutzomyia longipalpis*, and structurally characterise their *N*-glycan conjugates. We further discuss their implications for insect bloodfeeding as well as vector-host interactions.

## Results

### Identification of *Lutzomyia longipalpis* salivary glycoproteins

To determine the degree of *N*-glycosylation, an *in silico* analysis was carried out on 42 salivary proteins previously reported in *Lu. longipalpis*^4,20^ to predict protein *N*-glycosylation sites using the NetNGlyc server (http://www.cbs.dtu.dk/services/NetNGlyc/). This revealed 48% of the commonly known salivary proteins contain conventional *N*-glycosylation sites (Supplementary Table S1). However, it is important to note this list only includes proteins available on the NCBI database as studies published to date have focused on major secreted proteins, and no deep sequencing has been carried out for salivary glands of this sand fly species.

To accompany the *in silico* dataset, we carried out our own analysis of the sand fly salivary proteins (Supplementary Fig. S1). First, *Lu. longipalpis* salivary glands were dissected and individually pierced to release saliva. Subsequent Coomassie blue SDS-PAGE analysis showed several protein bands ranging from ∼10-100 kDa (Fig. 1). To identify which proteins were glycosylated, samples were analysed before and after treatment with Peptide-N-Glycosidase F (PNGase F), which cleaves high-mannose, hybrid and complex *N*-linked glycans. Treatment with PNGase F resulted in molecular mass shifts and migration of several protein bands, consistent with the widespread removal of *N*-glycans from the salivary glycoproteins (Fig. 1). De-glycosylation was also confirmed by transferring proteins to PVDF membrane and blotting with Concanavalin A (ConA) lectin, which binds specifically to terminal mannose residues on glycoproteins^21^ (Supplementary Fig. S2).

**Figure 1.**
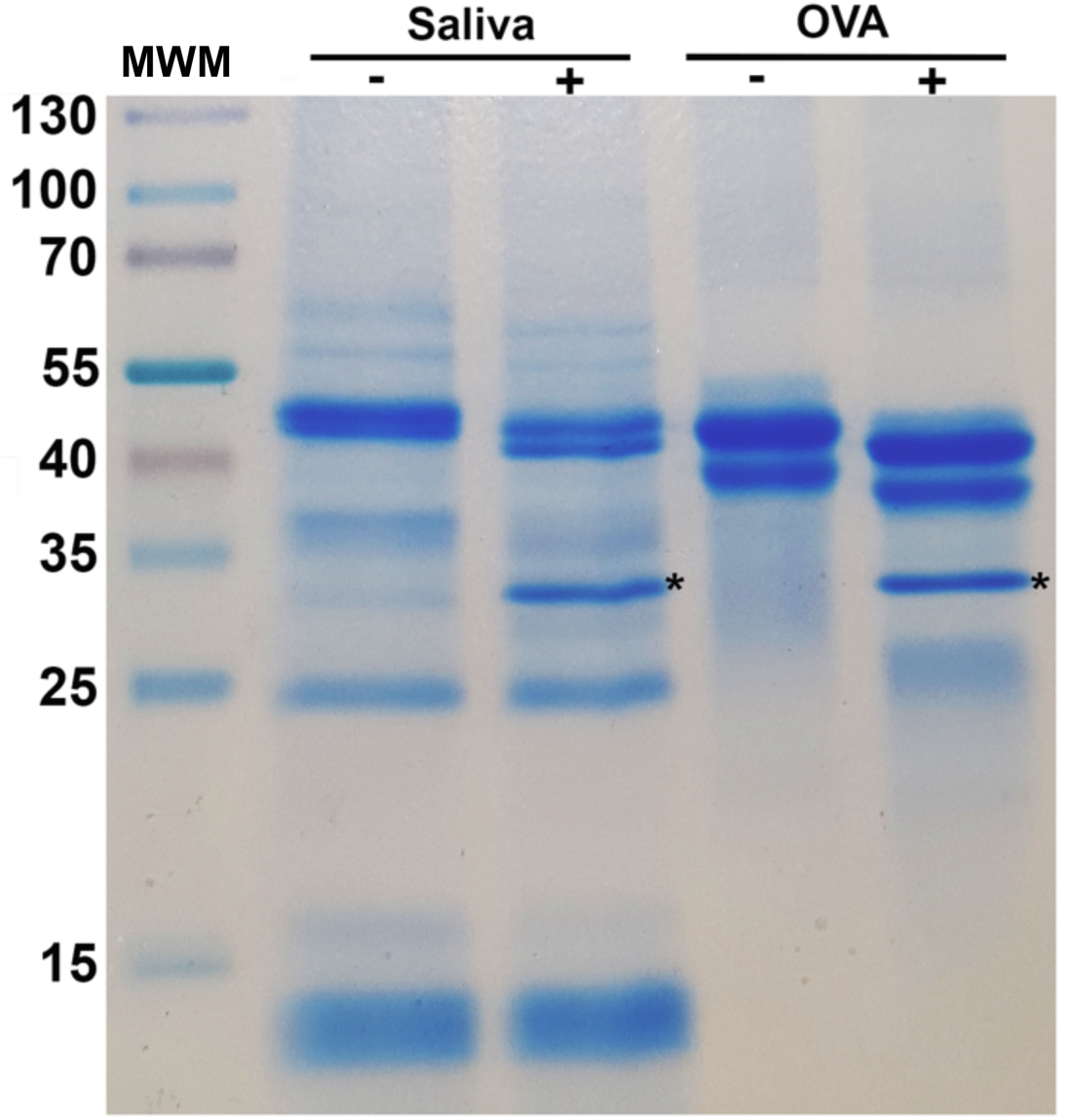
Enzymatic cleavage of *Lu. longipalpis* salivary glycoproteins with PNGase F. 10 µg of salivary proteins were incubated overnight with (+) and without (-) PNGase F to cleave *N*-glycans. Samples were resolved on a 12 % SDS-PAGE gel and Coomassie-stained. Egg albumin (OVA) was used as a positive control. MWM, molecular weight marker. *PNGase F enzyme.

For LC-MS/MS based glycoprotein identification, the major deglycosylated protein bands (Supplementary Fig. S3) were excised from the gel and sent to the University of Dundee Fingerprints Proteomics Facility. From the resulting list of 191 identified proteins, we excluded those without recognizable glycosylation sequons (as determined by NetNGlyc), obtaining a final list of 43 potentially *N*-glycosylated protein candidates (Supplementary Table S2). Fourteen of these potential glycoproteins were also identified in our initial *in silico* analysis (Supplementary Table S1), including LJM11, LJM111 and LJL143, which have been proposed as potential vaccine components against *Leishmania* infection^4^. Using the InterProScan tool to identify conserved protein domains, family distributions (Supplementary Fig. S4) show five of the candidates belonging to the actin family, while others like tubulin, 5’nucleotidase, peptidase M17 and the major royal jelly protein (yellow protein) are represented by two proteins each. After Blast2GO analysis, the “molecular function breakdown” suggested that 44% of the candidate glycoproteins are involved in binding, including ‘small molecule binding’ and ‘carbohydrate derivative binding’ (Supplementary Fig. S4). We also used the DeepLoc server to predict protein subcellular localisation and solubility of the proteins identified in Table S2. The results suggest 85% of candidate glycoproteins are soluble, and 10 proteins are both extracellular and soluble (Supplementary Table S2).

### Salivary glycoproteins from *Lu. longipalpis* are mainly modified with mannosylated *N*-glycans

Next, we determined the *N*-glycome modifying the salivary proteins of *Lu. longipalpis*. The presence of mannosylated *N*-glycan structures on salivary glycoproteins was suggested by the results of a lectin blot using Concanavalin A, and to confirm these results, we next determined the *N*-glycome of salivary glycoproteins of *Lu. longipalpis*.

The oligosaccharides were released by PNGase F followed by derivatization with procainamide^22^ which allowed fluorescence detection following hydrophilic interaction liquid chromatography (HILIC) and provided increased signal intensity in MS and MS/MS analysis^22^. Overall, we identified 14 different structures (Table 1), elucidated from ten separate compositions due to the presence of isomeric glycans.

Most oligosaccharides are of the high mannose type (82% of the *N*-glycome), with the Man_5_GlcNAc_2_-Proc glycan with *m/z* [727.81]^2+^, being the most abundant species (21.16 min; GU 6.00, Fig. 2). In addition, few hybrid-type species (with a retention time of 15.12-17.24 min) were detected, containing either an α1-6 core fucose residue linked to the reducing GlcNAc or not fucosylated, or a single terminal LacNAc motif (Fig. 2).

**Figure 2.**
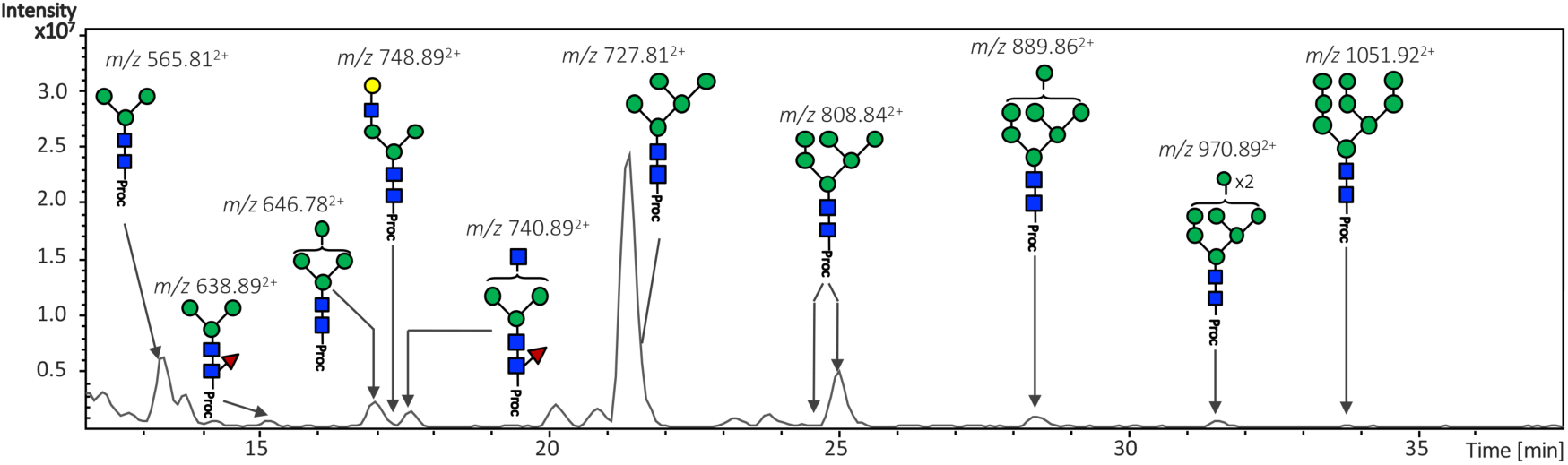
HILIC-LC separation of procainamide labelled *N*-glycans from *Lu. longipalpis*. Sand fly saliva contains mainly oligomannose-type *N*-linked glycans, with Man_5_GlcNAc_2_ being the most abundant structure. Green circle, mannose; yellow circle, galactose; Blue square, N-Acetylglucosamine; red triangle, fucose; Proc, procainamide.

All major glycan structures were characterised using positive ion MS (Fig. 3A) and MS/MS fragmentation spectra. An example of structural elucidation using MS/MS fragmentation spectrum is shown for the major glycan species Man_5_GlcNAc_2_-Proc, with *m/z* [727.82]^2+^ (Fig. 3B) while the remaining are mainly represented by hybrid-type glycans, either a trimannosyl modified with a Fuc residue on the chitobiose core, or paucimannosidic structures containing an unknown modification of 144 Da (see below).

**Figure 3.**
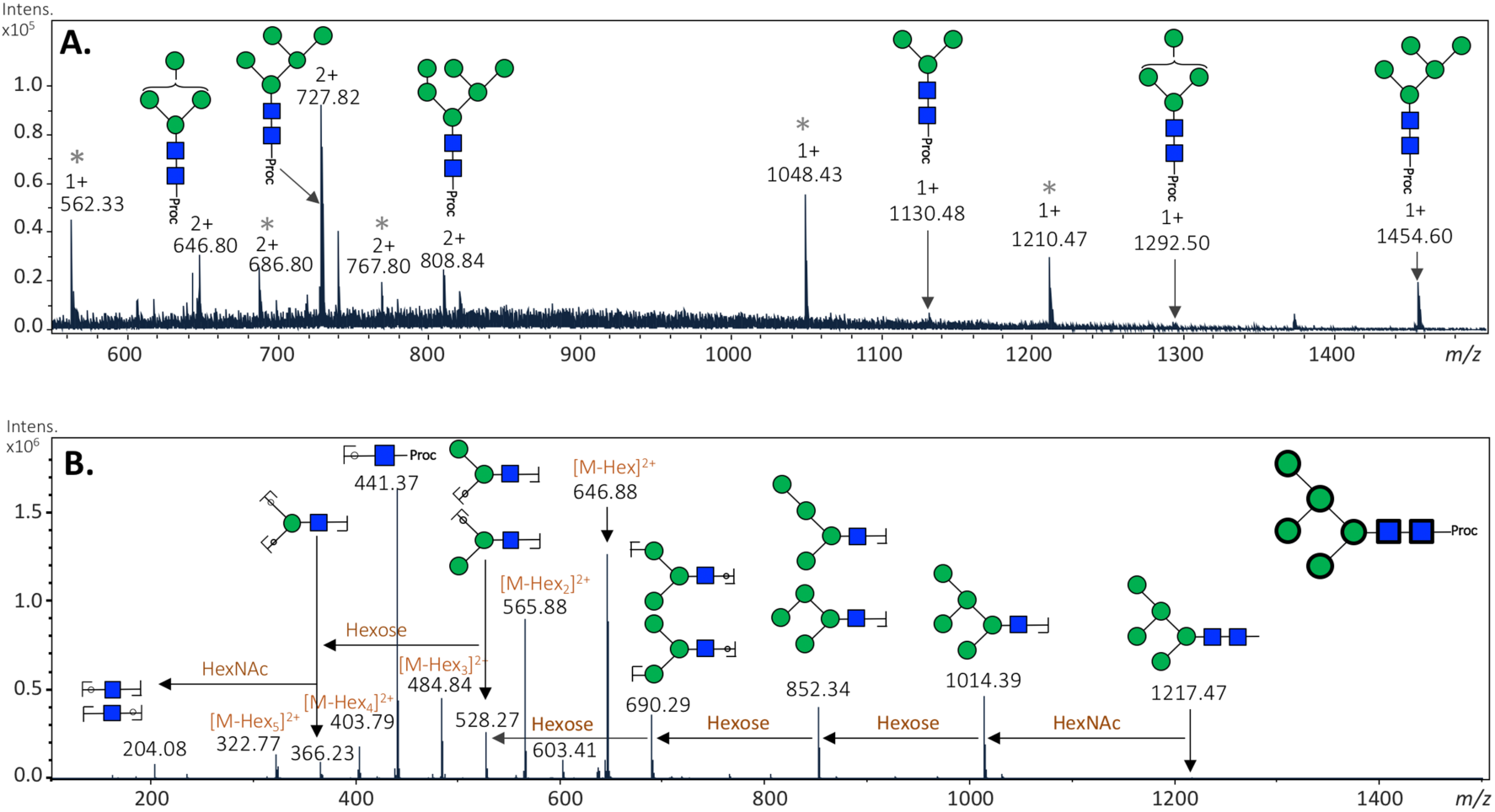
Mass spectrometry analysis of released *N*-glycans from *Lu. longipalpis* salivary glycoproteins. (A) Positive-ion mass spectrum profile (*m/z* 540-1,500) of total *N*-glycans. Ion signals are labelled accordingly. The most abundant glycan species (Hex_5_HexNAc_2_–Proc) was also detected as an [M+H]^2+^ ion with a *m/z* of 727.82. See Table 1 for complete glycan assignment. Peaks labelled with an asterisk correspond to glucose homopolymer contaminants from HILIC. (B) Positive-ion MS/MS fragmentation spectrum for most abundant *m/z* [727.8]^2+^ corresponding to the composition Hex_5_HexNAc_2_–Proc, proposed as a Man_5_GlcNAc_2_. Green circle, mannose; Blue square, N-Acetylglucosamine; Proc, procainamide.

Although PNGase F is highly effective in cleaving *N*-linked glycans, its activity is blocked by the presence of core fucose residues with an α1-3 linkage found in non-mammalian glycans. Therefore, we also treated our samples with PNGase A, which cleaves all glycans between the innermost GlcNAc and the asparagine independent of core linkages^23^. No differences were observed in chromatograms yielded from both enzymes (Supplementary Fig. S5), indicating all core fucosylation is likely to be α1-6-linked.

### MALDI-TOF-MS analysis reveals a series of sand fly salivary glycans with unidentified modifications of 144 Da

A more detailed analysis of the saliva by MALDI-TOF MS of pyridylaminated glycans revealed not only the major oligomannosidic species, but also suggested the existence of a series of glycans containing an unidentified structure. This modification was mainly found in two isomeric glycans: one with an RP-HPLC retention time of 25.0 min and the other of 26.5 min (Supplementary Fig. S6). The two isomers have a *m/z* 1295.50, which corresponds to a pyridylaminated Man_4_GlcNAc_2_ glycan carrying a modification of 144 Da. This was confirmed by treatment with Jack bean α-mannosidase, which resulted in a loss of 2 and 3 hexoses (Fig. 4) for each isomer, respectively. Interestingly, this modification seems to be located in different positions in the two structures, and in both cases this modification was lost after treatment with 48% aqueous hydrofluoric acid (aq.HF) (Fig. 4, and Table 2).

**Figure 4.**
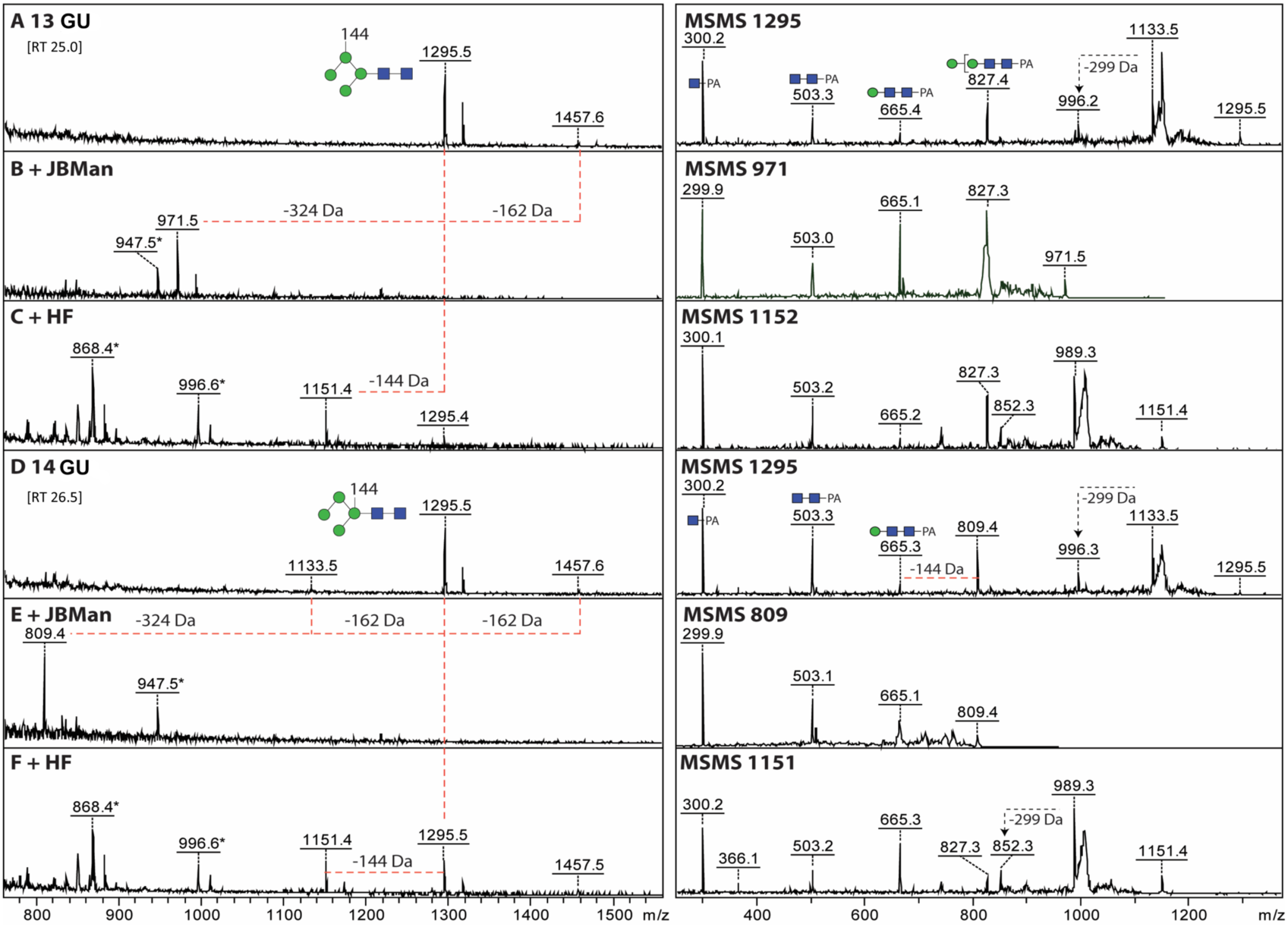
Analysis of sand fly *N*-glycans with an unknown residue. Two late-eluting RP-amide fractions (13 and 14 GU) containing glycans of *m/z* 1133, 1295 and 1457 (A, D) were analysed by MALDI-TOF MS and MS/MS before and after jack bean ⍰-mannosidase (B, E) or hydrofluoric acid (C, F) treatments. The *m/z* 1295 glycan structures lost either two or three mannose residues after mannosidase treatment, ruling out that terminal ⍰-mannose residues are substituted, but indicating a difference in the isomeric structure. In contrast, upon hydrofluoric acid treatment, incomplete loss of 144 Da was observed. Changes in mass upon mannosidase or HF treatment are indicated and non-glycan impurities annotated with an asterisk. The MS/MS for the original glycans and their digestion products are shown on the right; the differences in relative intensity of the *m/z* 665 and 827 fragments could explain the isomeric *m/z* 1295 structures with the 144 Da moiety attached to different mannose residues (as shown in panels A and D); key fragments are annotated according to the Symbolic Nomenclature for Glycans, while loss of reducing terminal GlcNAc-PA is indicated by -299 Da. PA, 2-aminopyridine; GU, glucose units; green circle, mannose; blue square, N-Acetylglucosamine.

Susceptibility to aq.HF is a hallmark of phosphoester, galactofuranose and some fucose modifications, but none of these are obviously compatible with a 144 Da modification. Based on this data, a re-assessment of the data with the procainamide-labelled glycans also revealed a potential oligosaccharide with a 144 Da modification (Supplementary Fig. S7); however, due to the very low abundance of these glycans we were unable to determine their chemical nature. Additionally, the potential for anionic modifications of *N*-glycans was explored by both glycomic workflows, but limitations in spectral quality and sample amount prevented a definitive characterisation.

### No *O*-linked glycans in sand fly saliva?

*In silico* predictions using the NetOGlyc 4.0^24^ server suggest that 85% our 191 identified salivary proteins have putative *O*-glycosylation sites (Supplementary Table S3). Sand fly saliva was subjected to reductive β-elimination to release *O*-glycans from the de-*N*-glycosylated proteins. Separation using porous graphitized carbon chromatography coupled with negative ion mode ESI-MS did not detect any *O*-glycans in the sample (Supplementary Fig. S8), either due to their absence, low abundance or low mass.

## Discussion

Sand fly saliva has important implications both for the insect and the vertebrate host^4^. *Lu. longipalpis* salivary proteins and their biological roles have been well studied^4,20^; however, the sugars that modify these proteins have not been characterised in detail. Most work on sand fly salivary glycans comes from *in silico* analyses^13-15,17,18,25^ and lectin blotting. They were first reported by Volf et al^19^, who used lectins to detect mannosylated *N*-type glycans. Mejia et al^16^ reported high mannose glycans in *Lu. longipalpis* saliva, with some potential hybrid-type structures (also based on lectin specificity). However, results from lectin-based methods should be interpreted with care as detection controls have not always been included in these studies, and results can be highly dependent on glycan abundance in samples and specific protocols. Our work is the first time that a mass spectrometry approach has been used to study the salivary *N*-linked glycans of *Lu. longipalpis*, providing detailed information about their structures and relative abundances. We found that sand fly salivary glycoproteins consist mainly of oligomannose glycans (ranging from the core Man_3_GlcNAc_2_ to Man_9_GlcNAc_2_), with some hybrid-type (e.g. fucosylated) structures. Additionally, this is the first report of a 144 Da (unknown) modification present in some salivary glycans. Our results provide new insights into how these structures could be recognised by vertebrate host cells.

In insects, protein glycosylation studies have been carried out primarily on the *Drosophila melanogaster* fly, demonstrating the presence of various carbohydrate structures^10,26,27^. It is generally accepted that *N*-linked type glycoproteins in arthropods are mainly of the high-mannose or paucimannose type, and account for over 90% of glycan complexity in *Drosophila*^10,28^. One of the first indications of the capacity of insects to produce complex type *N*-glycans came from bee venom phospholipase A2, which contains the core α1,3-fucose (an IgE epitope allergenic to humans). Anionic and zwitterionic *N*-glycans with up to three antennae have more recently been found in a range of insects^29-32^. Furthermore, Vandenborre et al.^33^ explored glycosylation differences comparing several economically important insects, and found glycoproteins to be involved in a broad range of biological processes such as cellular adhesion, homeostasis, communication and stress response.

Some researchers have predicted the presence of mucins in the mouthparts of bloodfeeders^34,35^, proposing their possible role as lubricants to facilitate bloodmeals. Even though *O*-linked glycans have been widely documented in invertebrates, we were unable to detect these sugars in sand fly saliva after reductive β-elimination. This was surprising given that our bioinformatic analysis (NetOGlyc server) predicted the presence of putative *O*-glycosylation sites. The presence of *O*-linked glycans in *Lu. longipalpis* saliva has been suggested through peanut agglutinin and *Vicia villosa* lectin detection^16^; however, it is worth noting that the experiment does not include positive controls or binding inhibition by competitive sugars, so non-specific binding cannot be ruled out. Interestingly, *Lu. longipalpis* midgut mucin-like glycoprotein has been described^36^ (with a suggested role in *Leishmania* attachment), showing the capacity of this species to produce *O*-linked glycans (at least in other tissues). A variety of *O*-linked glycans are reported for *Drosophila*^37^, with important functions such as body development ^10,38^. Furthermore, research shows that several *Drosophila*^37^ and moth^39^ cell lines form mucin-type *O*-glycans. It is worth noting there is no consensus sequence for *O*-glycosylation as in *N*-linked glycosylation, and *in silico* predictions are unreliable. Interestingly, similar results have been found in *Glossina* (unpublished), suggesting that these dipterans may not be able to *O*-glycosylate proteins in salivary tissues, or they are below the level of mass spectrometry detection.

A surprising finding in this work were the 144 Da structures modifying some of the salivary glycans (i.e. Man_4_GlcNAc_3_, and two Man_4_GlcNAc_2_ isomers). They were present in very low abundance (<1%), were located on different mannose residues (as shown by jack bean α-mannosidase digestion), and appeared susceptible to aqueous HF. However, we have yet to confirm the identity and biological role of this modification. A literature search revealed that structures of a 144 Da mass have been found on glycans from other organisms, including bacteria, viruses and sea algae^40-42^, but were not further addressed by the authors. One possibility is that these correspond to an anhydrosugar, like 3,6-anhydrogalactose (of 144 Da mass)^43^. Interestingly, work on mosquitoes has shown that these insects are able to produce anionic glycans with sulphate and/or glucuronic modifications that can be tissue specific^29,44^. The glycans identified here carrying this rare 144 Da residue may be another example of such modifications and could play a role specific to their location in sand fly saliva.

Even though every effort was made during salivary gland dissections to obtain saliva with minimal tissue contamination, this cannot be completely avoided. Analysis with the DeepLoc server suggested that although most protein candidates are ‘soluble’, only some are predicted to be ‘extracellular’. Furthermore, some proteins without signal peptide can still be secreted through a non-classical or “unconventional” secretory pathway^47,48^. An alternative way of saliva extraction would be to induce salivation by chemical means like pilocarpine^49-51^; however, this carries its own logistical difficulties considering the amount of saliva needed to detect glycans in such low abundances (even with the highly sensitive techniques we have used here). Another limitation of this work is the low protein profile resolution provided by 1D gel electrophoresis, where we may have missed weaker bands during our selection of proteins for sequencing. Higher protein concentrations and analysis through 2D gel electrophoresis could help us address this issue; nevertheless, we believe our work includes the major proteins in *Lu. longipalpis* saliva, providing a good overview of glycan abundance and composition in this bloodfeeding insect.

The biological role of protein glycosylation in the saliva of sand flies (and other bloodfeeding arthropods) is uncertain. One possibility is that glycans affect salivary protein half-life in the blood once they enter vertebrate host. Another possibility is that these glycans influence other *in vivo* processes like the interactions between saliva and cell surface carbohydrate recognition domains. For instance, the mannose receptor and DC-SIGN are c-type lectins that recognize mannosylated structures (uncommon in vertebrate cells); they are present on macrophages and dendritic cells, playing a role in both innate and adaptive immune systems^52^, making glycans highly relevant in parasitic infection processes. Additionally, the mannose-binding lectin activates the ‘lectin pathway’ of complement, and has an important role in protection against various pathogens^53^. An example of this was reported in tick saliva, which contains a mannose-binding lectin inhibitor whose activity was shown to be glycosylation-dependent^54^.

This, in turn, could be of importance within the context of *Leishmania* infection as both macrophages and dendritic cells have been shown to have critical roles in the initial stages of infection and subsequent dissemination of the parasite inside the vertebrate host^55^. In order for *Leishmania* to survive and multiply inside the host, it must be internalized by macrophages; however, promastigotes appear to avoid the MR receptor during invasion, as it promotes inflammation and can be detrimental to their survival^55^. The saliva of *Lu. longipalpis* can prevent macrophages from presenting *Leishmania* antigens to T cells^56^, but these effects are species-specific; in the case of other sand flies like *Phlebotomus papatasi*, saliva inhibits the activation of these cells^57^. Work on a patient-isolated *L. major* strain that causes nonhealing lesions in C57BL/6 mice found that its uptake by dermal-macrophages is MR-mediated^58^. Even though the MR does not play a role in the healing strain, it is an indication that sand fly saliva may be involved in other parasite-macrophage interactions. *Leishmania* also interacts with DC-SIGN (particularly amastigotes and metacyclic promastigotes) and this varies depending on species^59^. It remains to be seen whether mannosylated glycoproteins in saliva impair or facilitate these interactions and their outcomes.

Many sand fly salivary proteins are currently being explored as potential vaccine candidates against *Leishmania*, and knowing the nature of their post-translational modifications is relevant to their activity and efficacy. Several salivary proteins from *Lu. longipalpis* that are being researched as vaccine candidates (e.g. LJM11, LJM17 and LJL143^4^) have potential glycosylation sites (as indicated in the results of our *in silico* analysis). As recombinant versions of these proteins are normally expressed in non-insect cells^60^, care should be taken to ensure the glycoprotein’s profile and activity remains the same.

Finally, it is also worth considering the role salivary glycoproteins could play inside the sand flies themselves. Both male and female sand flies rely on plant sugars to survive, and Cavalcante et al. showed that *Lu. longipalpis* ingest saliva while sugar feeding^61^. Lectins (which bind to glycans) represent a major part of a plant’s defence system^62^, and can cause damage to an insect’s midgut when ingested^63^. Salivary glycoconjugates may be potentially recognized by these plant lectins, helping to decrease the damage they can cause. Moreover, the ingestion of saliva during the bloodmeal may impact parasite differentiation in the fly’s gut^64^. Furthermore, sand fly-borne viruses use the host cell machinery for replication, which includes the insect glycosylation pathways, before it is transmitted to the vertebrate host. In this context, understanding the glycosylation of insect salivary glands is also relevant to understand their pathogenicity.

## Methods

### Glycoprotein predictions

The servers NetNGlyc 1.0^65^ (http://www.cbs.dtu.dk/services/NetNGlyc/) and NetOGlyc 4.0^24,66^ (http://www.cbs.dtu.dk/services/NetOGlyc/) were used to predict potential glycosylation sites by examination of the consensus sequences. The DeepLoc 1.0^67^ server (http://www.cbs.dtu.dk/services/DeepLoc/index.php) was used to predict location of proteins.

### Sand fly salivary gland dissection and extraction of saliva

*Lutzomyia longipalpis* sand flies were obtained from a colony at the London School of Hygiene and Tropical Medicine (UK), which originated in Jacobina (Bahia state), Brazil. Salivary glands were dissected from 5-day old, sugar-fed, uninfected females in sterile PBS (Sigma, St. Louis, US). To harvest saliva, pools of 10 salivary glands were placed on ice, pierced with a needle and then centrifuged at 3000 rpm for 10 min at 4°C. The supernatant (pure saliva) was stored at -80°C. Between 0.5-1 μg of protein per sand fly was obtained from dissections.

### SDS polyacrylamide gel electrophoresis and staining

Sand fly saliva (10 μg) was run on a 12.5% polyacrylamide gel, before and after deglycosylation with endoglycosidase PNGase F (New England Biolabs, Massachusetts, US). Gel was stained using InstantBlue Protein stain (Expedeon, California, US). Spectra Multicolor Broad Range Protein Ladder (ThermoFisher, UK) was used as molecular weight marker.

### Concanavalin A blots

Saliva samples, before and after treatment with PNGase F (New England Biolabs, US) were run on a 12.5% polyacrylamide gel under standard conditions, transferred onto a PVDF membrane (Fisher Scientific, UK), and blocked with 1% BSA (Sigma, St. Louis, US) in PBS-Tw 20 (Sigma, St. Louis, US) overnight at 4°C. Membrane was incubated with 1 μg/ml biotinylated Concanavalin A (ConA) lectin (Vector Labs, Peterborough, UK) for 1 hour at room temperature. After washing, the membrane was incubated with 1:100,000 streptavidin-HRP (Vector Labs, Peterborough, UK). SuperSignal West Pico Chemiluminescent substrate (ThermoFisher, Massachusetts, US) was used to detect the bands. Egg albumin (Sigma, St. Louis, US), a highly mannosylated *N*-linked glycoprotein^68^, was used as positive control.

### Mass spectrometry analysis

To identify the glycoproteins that were susceptible to PNGase F, bands of interest were sliced from the gel and sent to the Dundee University Fingerprints Proteomics Facility. Briefly, the excised bands were subjected to in-gel trypsination then alkylated with iodoacetamide. The resultant peptides were then analysed via liquid chromatography-tandem mass spectrometry (LC-MS/MS) in a Thermo LTQ XL Linear Trap instrument equipped with a nano-LC. Tandem MS data were searched against the *Lu. longipalpis* database downloaded from VectorBase (https://www.vectorbase.org/proteomes) using the Mascot (version 2.3.02, Matrix Science, Liverpool) search engine. Search parameters were performed as described in elsewhere^69^. For in-solution data, the false discovery rate was filtered at 1%, and individual ion scores ≥30 were considered to indicate identity or extensive homology (p<0.05).

### Enzymatic release of *N*-linked glycans

The *N*-glycans from sand fly saliva were released by in-gel deglycosylation using PNGase F as described by Royle *et al*.^70^. For deglycosylation using PNGase A, peptides were released from gel pieces by overnight incubation at 37 °C with trypsin in 25 mM ammonium bicarbonate. The supernatant was dried, re-suspended in water and heated at 100 °C for 10 min to deactivate the trypsin. Samples were dried by vacuum centrifugation and the tryptic peptide mixture was incubated with PNGase A in 100 mM citrate/phosphate buffer (pH 5.0) for 16 h at 37 °C^71^. Samples were separated from protein and salts using LudgerClean Protein Binding Plate (Ludger Ltd., Oxfordshire, UK). All wells were flushed with extra water to ensure full recovery and then dried by vacuum centrifugation prior to fluorescent labelling.

### Fluorescent labelling and purification of released *N*-glycans

Released *N*-glycans were fluorescently labelled via reductive amination reaction with procainamide using a Ludger Procainamide Glycan Labelling Kit containing 2-picoline borane (Ludger Ltd.). The released glycans were incubated with labelling reagents for 1 h at 65 °C. The procainamide labelled glycans were cleaned up using LudgerClean S Cartridges (Ludger Ltd) and eluted with water (1 mL). Samples were evaporated under high vacuum and re-suspended in water prior to use.

### ESI-LC-MS and ESI-LC-MS/MS analysis of procainamide-labelled *N*-glycans

Procainamide labelled samples were analysed by ESI-LC-MS in positive ion mode. 25 µL of each sample were injected onto an ACQUITY UPLC BEH-Glycan 1.7 µm, 2.1 × 150 mm column at 40 °C on the Dionex Ultimate 3000 UHPLC attached to a Bruker Amazon Speed ETD (Bruker, UK). The running conditions used were: solvent A was 50 mM ammonium formate pH 4.4; solvent B was acetonitrile (acetonitrile 190 far UV/gradient quality; Romil #H049). Gradient conditions were: 0 to 53.5 min, 24% A (0.4 mL/min); 53.5 to 55.5 min, 24 to 49 % A (0.4 mL/min); 55.5 to 57.5min, 49 to 60% A (0.4 to 0.25 mL/min); 57.5 to 59.5 min, 60% A (0.25 mL/min); 59.5 to 65.5 min, 60 to 24% A (0.4 mL/min); 65.5 to 66.5 min, 24% A (0.25 to 0.4 mL/min); 66.5 to 70 min 24% A (0.4 mL/min). The Amazon Speed settings were the same as described in^72^ except that precursor ions were released after 0.2 min and scanned in enhanced resolution within a mass range of 200-1500 *m/z* (target mass, 900 *m/z*).

### Release of *O*-linked glycans

Saliva samples underwent reductive β-elimination to release *O*-glycans after PNGase F treatment. Briefly, samples were diluted in 0.05 M sodium hydroxide and 1.0 M sodium borohydride at a temperature of 45°C with an incubation time of 14-16 h followed by solid-phase extraction of released *O*-glycans^73^. *O*-glycans were analysed using PGC-LC coupled to negative ion ESI-MS/MS^74^ alongside bovine fetuin *O*-glycans as a positive control.

### MALDI-TOF analysis of aminopyridine-labelled glycans

Sand fly salivary glycans were released according to previous procedures and labelled with PA (aminopyridine) as described elsewhere^75^, prior to RP-HPLC and analysis by MALDI-TOF MS using a Bruker Daltonics Autoflex Speed instrument (Hykollari). Aliquots of samples were treated with Jack bean α-mannosidase (Sigma), α-1,3 mannosidase and 48% aqueous hydrofluoric acid (aq.HF); the latter under control conditions releases phospho(di)esters, phosphonate, α1,3-fucose and galactofuranose groups. Dried glycan fractions were redissolved in 3 μL of aq.HF on ice (in the cold room) for 36 h prior to repeated evaporation. The digests were re-analysed using MALDI-TOF MS and MS/MS. Spectra were annotated by comparison to previous data on insect N-glycomes in terms of monosaccharide composition (Fx Hy Nz), using retention time, manual interpretation, exoglycosidase treatment results and LIFT fragmentation analysis.

## Acknowledgements

This work was supported in part by a Ph.D. studentship by the Colombian Department of Science, Technology and Innovation (Colciencias) through the scholarship programme “Francisco José de Caldas” (to KMS) and by the GlycoPar Marie Curie Initial Training Network GA 608295 (to KWK, DS, IW and AA-S). The Biotechnology and Biological Sciences Research Council supported MER through a David Phillips Fellowship (BB/H022406/1). The funders had no role in study design, data collection and analysis, decision to publish, or preparation of the manuscript. We thank Douglas Lamont (Dundee University Fingerprints Facility) for assistance with proteomics identification of sand fly salivary proteins.

## Author contributions

Designed experiments (KMS, KWK, DS, AA-S), performed experiments (KMS, KWK, SY, RK) and analysed the data (KMS, KWK, SY, IW, KP, RK, MER, AA-S), wrote the manuscript (KMS, KWK, AA-S). All authors reviewed and approved the manuscript.

## TABLES

**Table 1.** List of glycan structures present in *Lu. longipalpis* saliva. GU, glucose unit; Proc, procainamide. Green circles, mannose; Blue squares, N-Acetylglucosamine; Red triangle, fucose; yellow circles, galactose.

**Table 2.** Summary of treatments of the isomeric structures detected by MALDI-TOF-MS (Fig 4). JBMan, Jack Bean α-mannosidase; GU, glucose units; RT, retention time; aq.HF, aqueous Hydrofluoric acid.

## References

1 Maroli, M., Feliciangeli, M. D., Bichaud, L., Charrel, R. N. & Gradoni, L. Phlebotomine sandflies and the spreading of leishmaniases and other diseases of public health concern. Med Vet Entomol 27, 123–147, doi: 10.1111/j.1365-2915.2012.01034.x (2013).

2 Staudacher, E. et al. Alpha 1-6(alpha 1-3)-difucosylation of the asparagine-bound *N*-acetylglucosamine in honeybee venom phospholipase A2. Glycoconj J 9, 82–85 (1992).

3 World Health Organization. Control of the leishmaniasis: report of a meeting of the WHO Expert Committee on the Control of the Leishmaniasis. Report No. 949, (Geneva, 2010).

4 Abdeladhim, M., Kamhawi, S. & Valenzuela, J. G. What’s behind a sand fly bite? The profound effect of sand fly saliva on host hemostasis, inflammation and immunity. Infect Genet Evol 28, 691–703, doi: 10.1016/j.meegid.2014.07.028 (2014).

5 Lestinova, T., Rohousova, I., Sima, M., de Oliveira, C. I. & Volf, P. Insights into the sand fly saliva: Blood-feeding and immune interactions between sand flies, hosts, and *Leishmania*. PLoS Negl Trop Dis 11, e0005600, doi: 10.1371/journal.pntd.0005600 (2017).

6 Rogers, M. E. The role of leishmania proteophosphoglycans in sand fly transmission and infection of the Mammalian host. Front Microbiol 3, 223, doi: 10.3389/fmicb.2012.00223 (2012).

7 Mondragon-Shem, K. et al. Severity of old world cutaneous leishmaniasis is influenced by previous exposure to sandfly bites in Saudi Arabia. PLoS Negl Trop Dis 9, e0003449, doi: 10.1371/journal.pntd.0003449 (2015).

8 Cecilio, P. et al. Pre-clinical antigenicity studies of an innovative multivalent vaccine for human visceral leishmaniasis. PLoS Negl Trop Dis 11, e0005951, doi: 10.1371/journal.pntd.0005951 (2017).

9 Rudd, P., Elliott, T., Cresswell, P., Wilson, I. & Dwek, R. Glycosylation and the immune system. Science 291 (2001).

10 Katoh, T. & Tiemeyer, M. The N’s and O’s of *Drosophila* glycoprotein glycobiology. Glycoconj J 30, 57–66, doi: 10.1007/s10719-012-9442-x (2013).

11 Rogers, M. E., Ilg, T., Nikolaev, A. V., Ferguson, M. A. & Bates, P. A. Transmission of cutaneous leishmaniasis by sand flies is enhanced by regurgitation of fPPG. Nature 430, 463–467, doi: 10.1038/nature02675 (2004).

12 Gomes, R. & Oliveira, F. The immune response to sand fly salivary proteins and its influence on *Leishmania* immunity. Front Immunol 3 (2012).

13 Abdeladhim, M. et al. Updating the salivary gland transcriptome of *Phlebotomus papatasi* (Tunisian strain): the search for sand fly-secreted immunogenic proteins for humans. PLoS One 7, e47347, doi: 10.1371/journal.pone.0047347 (2012).

14 Hostomska, J. et al. Analysis of salivary transcripts and antigens of the sand fly *Phlebotomus arabicus*. BMC Genomics 10, 282, doi: 10.1186/1471-2164-10-282 (2009).

15 Martin-Martin, I., Molina, R. & Jimenez, M. Identifying salivary antigens of *Phlebotomus argentipes* by a 2DE approach. Acta Trop 126, 229–239, doi: 10.1016/j.actatropica.2013.02.008 (2013).

16 Mejia, J. S., Toot-Zimmer, A. L., Schultheiss, P. C., Beaty, B. J. & Titus, R. G. BluePort: a platform to study the eosinophilic response of mice to the bite of a vector of *Leishmania* parasites, *Lutzomyia longipalpis* sand flies. PLoS One 5, e13546, doi: 10.1371/journal.pone.0013546 (2010).

17 Rohousova, I. et al. Salivary gland transcriptomes and proteomes of *Phlebotomus tobbi* and *Phlebotomus sergenti*, vectors of leishmaniasis. PLoS Negl Trop Dis 6, e1660, doi: 10.1371/journal.pntd.0001660 (2012).

18 Vlkova, M. et al. Comparative analysis of salivary gland transcriptomes of *Phlebotomus orientalis* sand flies from endemic and non-endemic foci of visceral leishmaniasis. PLoS Negl Trop Dis 8, e2709, doi: 10.1371/journal.pntd.0002709 (2014).

19 Volf, P., Tesarova, P. & Nohynkova, E. N. Salivary proteins and glycoproteins in phlebotomine sandflies of various species, sex and age. Med Vet Entomol 14, 251–256 (2000).

20 Valenzuela, J. G., Garfield, M., Rowton, E. D. & Pham, V. M. Identification of the most abundant secreted proteins from the salivary glands of the sand fly *Lutzomyia longipalpis*, vector of *Leishmania chagasi*. J Exp Biol 207, 3717–3729, doi: 10.1242/jeb.01185 (2004).

21 Maupin, K. A., Liden, D. & Haab, B. B. The fine specificity of mannose-binding and galactose-binding lectins revealed using outlier motif analysis of glycan array data. Glycobiology 22, 160–169, doi: 10.1093/glycob/cwr128 (2012).

22 Kozak, R. P., Tortosa, C. B., Fernandes, D. L. & Spencer, D. I. Comparison of procainamide and 2-aminobenzamide labeling for profiling and identification of glycans by liquid chromatography with fluorescence detection coupled to electrospray ionization-mass spectrometry. Anal Biochem 486, 38–40, doi: 10.1016/j.ab.2015.06.006 (2015).

23 Wang, T. et al. Discovery and characterization of a novel extremely acidic bacterial *N*-glycanase with combined advantages of PNGase F and A. Biosci Rep 34, e00149, doi: 10.1042/BSR20140148 (2014).

24 Hansen, J. E. et al. NetOglyc: prediction of mucin type O-glycosylation sites based on sequence context and surface accessibility. Glycoconj J 15, 115–130 (1998).

25 Sima, M. et al. The Diversity of Yellow-Related Proteins in Sand Flies (Diptera: Psychodidae). PLoS One 11, e0166191, doi: 10.1371/journal.pone.0166191 (2016).

26 Seppo, A. & Tiemeyer, M. Function and structure of *Drosophila* glycans. Glycobiology 10, 751–760, doi: 10.1093/glycob/10.8.751 (2000).

27 ten Hagen, K. G., Zhang, L., Tian, E. & Zhang, Y. Glycobiology on the fly: developmental and mechanistic insights from *Drosophila*. Glycobiology 19, 102–111, doi: 10.1093/glycob/cwn096 (2009).

28 Aoki, K. et al. Dynamic developmental elaboration of *N*-linked glycan complexity in the *Drosophila melanogaster* embryo. J Biol Chem 282, 9127–9142, doi: 10.1074/jbc.M606711200 (2007).

29 Kurz, S. et al. Targeted release and fractionation reveal glucuronylated and sulphated N-and O-glycans in larvae of dipteran insects. J Proteomics 126, 172–188, doi: 10.1016/j.jprot.2015.05.030 (2015).

30 Cabrera, G. et al. Structural characterization and biological implications of sulfated *N*-glycans in a serine protease from the neotropical moth *Hylesia metabus* (Cramer [1775]) (Lepidoptera: Saturniidae). Glycobiology 26, 230–250, doi: 10.1093/glycob/cwv096 (2016).

31 Hykollari, A. et al. Isomeric separation and recognition of anionic and zwitterionic *N*-glycans from royal jelly glycoproteins. Mol Cell Proteomics 17, 2177–2196, doi: 10.1074/mcp.RA117.000462 (2018).

32 Stanton, R. et al. The underestimated N-glycomes of lepidopteran species. Biochim Biophys Acta Gen Subj 1861, 699–714, doi: 10.1016/j.bbagen.2017.01.009 (2017).

33 Vandenborre, G. et al. Diversity in protein glycosylation among insect species. PloS one 6, e16682, doi: 10.1371/journal.pone.0016682 (2011).

34 Francischetti, I. M., Valenzuela, J. G., Pham, V. M., Garfield, M. K. & Ribeiro, J. M. Toward a catalog for the transcripts and proteins (sialome) from the salivary gland of the malaria vector *Anopheles gambiae*. J Exp Biol 205, 2429–2451 (2002).

35 Esteves, E. et al. Analysis of the salivary gland transcriptome of unfed and partially fed *Amblyomma sculptum* ticks and descriptive proteome of the saliva. Front Cell Infect Microbiol 7, 476, doi: 10.3389/fcimb.2017.00476 (2017).

36 Myskova, J. et al. Characterization of a midgut mucin-like glycoconjugate of *Lutzomyia longipalpis* with a potential role in *Leishmania* attachment. Parasit Vectors 9, 413, doi: 10.1186/s13071-016-1695-y (2016).

37 Tiemeyer, M., Nakato, H. & Esko, J. D. in Essentials of Glycobiology (eds rd et al.) 335–349 (2015).

38 Nakamura, N., Lyalin, D. & Panin, V. M. Protein O-mannosylation in animal development and physiology: from human disorders to *Drosophila* phenotypes. Semin Cell Dev Biol 21, 622–630, doi: 10.1016/j.semcdb.2010.03.010 (2010).

39 Staudacher, E. Mucin-Type O-Glycosylation in Invertebrates. Molecules 20, 10622–10640, doi: 10.3390/molecules200610622 (2015).

40 Piacente, F. et al. Giant DNA virus mimivirus encodes pathway for biosynthesis of unusual sugar 4-amino-4,6-dideoxy-D-glucose (Viosamine). J Biol Chem 287, 3009–3018, doi: 10.1074/jbc.M111.314559 (2012).

41 Thomas, R. M. et al. Glycosylation of DsbA in *Francisella tularensis* subsp. tularensis. J Bacteriol 193, 5498–5509, doi: 10.1128/JB.00438-11 (2011).

42 Black, G. E., Fox, A., Fox, K., Snyder, A. P. & Smith, P. B. Electrospray tandem mass spectrometry for analysis of native muramic acid in whole bacterial cell hydrolysates. Anal Chem 66, 4171–4176, doi: 10.1021/ac00095a010 (1994).

43 Yaphe, W. Colorimetric Determination of 3,6-Anhydrogalactose with the Indolyl-3-acetic Acid Reagent. Nature 197, 488–489 (1963).

44 Hykollari, A., Malzl, D., Stanton, R., Eckmair, B. & Paschinger, K. Tissue-specific glycosylation in the honeybee: Analysis of the N-glycomes of *Apis mellifera* larvae and venom. Biochim Biophys Acta Gen Subj 1863, 129409, doi: 10.1016/j.bbagen.2019.08.002 (2019).

45 Valenzuela, J. G., Belkaid, Y., Rowton, E. & Ribeiro, J. M. The salivary apyrase of the blood-sucking sand fly *Phlebotomus papatasi* belongs to the novel *Cimex* family of apyrases. J Exp Biol 204, 229–237 (2001).

46 Charlab, R., Valenzuela, J. G., Rowton, E. D. & Ribeiro, J. M. Toward an understanding of the biochemical and pharmacological complexity of the saliva of a hematophagous sand fly *Lutzomyia longipalpis*. Proc Natl Acad Sci U S A 96, 15155–15160, doi: 10.1073/pnas.96.26.15155 (1999).

47 Rabouille, C. Pathways of Unconventional Protein Secretion. Trends Cell Biol 27, 230–240, doi: 10.1016/j.tcb.2016.11.007 (2017).

48 Nickel, W. & Rabouille, C. Mechanisms of regulated unconventional protein secretion. Nat Rev Mol Cell Biol 10, 148–155, doi: 10.1038/nrm2617 (2009).

49 Ribeiro, J. M., Zeidner, N. S., Ledin, K., Dolan, M. C. & Mather, T. N. How much pilocarpine contaminates pilocarpine-induced tick saliva? Med Vet Entomol 18, 20–24, doi: 10.1111/j.0269-283x.2003.0469.x (2004).

50 Oliveira, C. J. et al. Proteome of *Rhipicephalus sanguineus* tick saliva induced by the secretagogues pilocarpine and dopamine. Ticks Tick Borne Dis 4, 469–477, doi: 10.1016/j.ttbdis.2013.05.001 (2013).

51 Boorman, J. Induction of salivation in biting midges and mosquitoes, and demonstration of virus in the saliva of infected insects. Med Vet Entomol 1, 211–214, doi: 10.1111/j.1365-2915.1987.tb00346.x (1987).

52 Taylor, P. R. et al. Macrophage receptors and immune recognition. Annu Rev Immunol 23, 901–944, doi: 10.1146/annurev.immunol.23.021704.115816 (2005).

53 Schnaar, R. L. Glycobiology simplified: diverse roles of glycan recognition in inflammation. J Leukoc Biol 99, 825–838, doi: 10.1189/jlb.3RI0116-021R (2016).

54 Schuijt, T. J. et al. A tick mannose-binding lectin inhibitor interferes with the vertebrate complement cascade to enhance transmission of the lyme disease agent. Cell Host Microbe 10, 136–146, doi: 10.1016/j.chom.2011.06.010 (2011).

55 Liu, D. & Uzonna, J. The early interaction of *Leishmania* with macrophages and dendritic cells and its influence on the host immune response. Front Cell Infect Microbiol 2 (2012).

56 Theodos, C. M. & Titus, R. G. Salivary-Gland Material from the Sand Fly Lutzomyia-Longipalpis Has an Inhibitory Effect on Macrophage Function in-Vitro. Parasite Immunology 15, 481–487, doi: DOI 10.1111/j.1365-3024.1993.tb00634.x (1993).

57 Hall, L. R. & Titus, R. G. Sand fly vector saliva selectively modulates macrohage functions that inhibit killing of *Leishmania* production. The Journal of Immunology 155, 3501–3506 (1995).

58 Lee, S. H. et al. Mannose receptor high, M2 dermal macrophages mediate nonhealing *Leishmania major* infection in a Th1 immune environment. J Exp Med 215, 357–375, doi: 10.1084/jem.20171389 (2018).

59 Caparros, E. et al. Role of the C-type lectins DC-SIGN and L-SIGN in *Leishmania* interaction with host phagocytes. Immunobiology 210, 185–193, doi: 10.1016/j.imbio.2005.05.013 (2005).

60 Hamasaki, R., Kato, H., Terayama, Y., Iwata, H. & Valenzuela, J. G. Functional characterization of a salivary apyrase from the sand fly, Phlebotomus duboscqi, a vector of Leishmania major. J Insect Physiol 55, 1044–1049, doi: 10.1016/j.jinsphys.2009.07.010 (2009).

61 Cavalcante, R., Pereira, M., Freitas, J. & de F Gontijo, N. Ingestion of saliva during carbohydrate feeding by *Lutzomyia longipalpis* (Diptera; Psychodidae). Mem Inst Oswaldo Cruz 10 (2006).

62 Lannoo, N. & Van Damme, E. J. Lectin domains at the frontiers of plant defense. Front Plant Sci 5, 397, doi: 10.3389/fpls.2014.00397 (2014).

63 Zhu-Salzman, K. & Zeng, R. Insect response to plant defensive protease inhibitors. Annu Rev Entomol 60, 233–252, doi: 10.1146/annurev-ento-010814-020816 (2015).

64 Charlab, R. & Ribeiro, J. M. Cytostatic efect of *Lutzomyia longipalpis* salivary gland homogenates on *Leishmania* parasites. Am J Trop Med Hyg 48, 831–838 (1993).

65 Gupta, R., Jung, E. & Brunak, S. NetNGlyc 1.0 Server, <http://www.cbs.dtu.dk/services/NetNGlyc/> (2017).

66 Steentoft, C. et al. (DTU Bioinformatics. Department of Bio and Health Informatics, 2017).

67 Almagro Armenteros, J. J., Sonderby, C. K., Sonderby, S. K., Nielsen, H. & Winther, O. DeepLoc: prediction of protein subcellular localization using deep learning. Bioinformatics 33, 3387–3395, doi: 10.1093/bioinformatics/btx431 (2017).

68 Harvey, D. J., Wing, D. R., Kuster, B. & Wilson, I. B. Composition of N-linked carbohydrates from ovalbumin and co-purified glycoproteins. J Am Soc Mass Spectrom 11, 564–571, doi: 10.1016/S1044-0305(00)00122-7 (2000).

69 Rose, C. et al. An investigation into the protein composition of the teneral *Glossina morsitans morsitans* peritrophic matrix. PLoS Negl Trop Dis 8, e2691, doi: 10.1371/journal.pntd.0002691 (2014).

70 Royle, L., Radcliffe, C. M., Dwek, R. A. & Rudd, P. M. Detailed structural analysis of N-glycans released from glycoproteins in SDS-PAGE gel bands using HPLC combined with exoglycosidase array digestions. Methods Mol Biol 347, 125–143, doi: 10.1385/1-59745-167-3:125 (2006).

71 Navazio, L. et al. Monitoring endoplasmic reticulum-to-Golgi traffic of a plant calreticulin by protein glycosylation analysis. Biochemistry 41, 14141–14149, doi: 10.1021/bi0204701 (2002).

72 Kotsias, M. et al. Method comparison for N-glycan profiling: Towards the standardization of glycoanalytical technologies for cell line analysis. PLoS One 14, e0223270, doi: 10.1371/journal.pone.0223270 (2019).

73 Carlson, D. M. Structures and immunochemical properties of oligosaccharides isolated from pig submaxillary mucins. J Biol Chem 243, 616–626 (1968).

74 Jensen, P. H., Karlsson, N. G., Kolarich, D. & Packer, N. H. Structural analysis of N- and O-glycans released from glycoproteins. Nat Protoc 7, 1299–1310, doi: 10.1038/nprot.2012.063 (2012).

75 Paschinger, K. et al. The N-glycans of *Trichomonas vaginalis* contain variable core and antennal modifications. Glycobiology 22, 300–313, doi: 10.1093/glycob/cwr149 (2012).

