## Supplementary Figures for "Insights into the salivary *N*-glycome of *Lutzomyia longipalpis*, vector of visceral leishmaniasis"

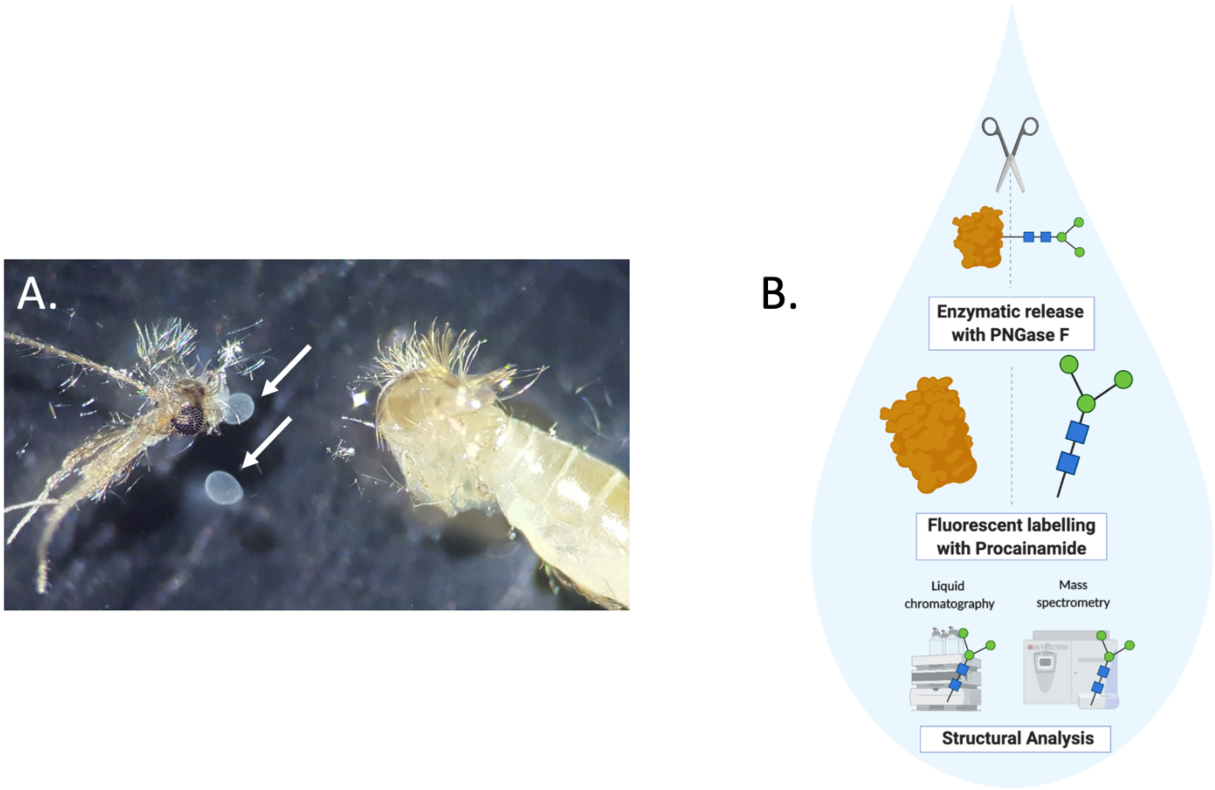


**Supplementary Figure S1. Analysis of the sandfly glycosialome.** A. Saliva was dissected from 5-day old, sugar fed Lu. longipalpis females (white arrows indicated extracted salivary glands). B. Glycoproteins were subjected to enzymatic release by PNGase F, and glycans were fluorescently labelled with procainamide for analysis by LC and MS. Figure created with BioRender.

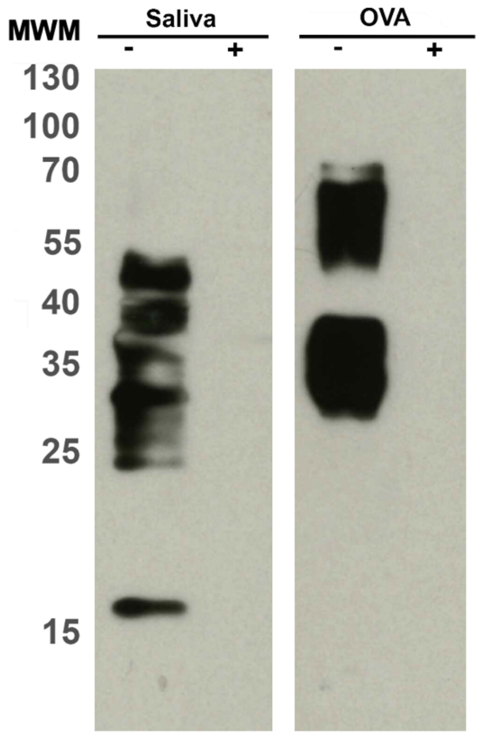


Supplementary Figure S2. Detection of *Lu. longipalpis* saliva mannosylated glycoproteins with Concanavalin A. 10 µg of salivary proteins were incubated overnight with (+) and without (-) PNGase F to cleave *N*-glycans. Samples were resolved on a 12 % SDS-PAGE gel and transferred onto a PVDF membrane and blotted with Con A. Egg albumin (OVA) was used as a positive control. MWM: molecular weight marker.

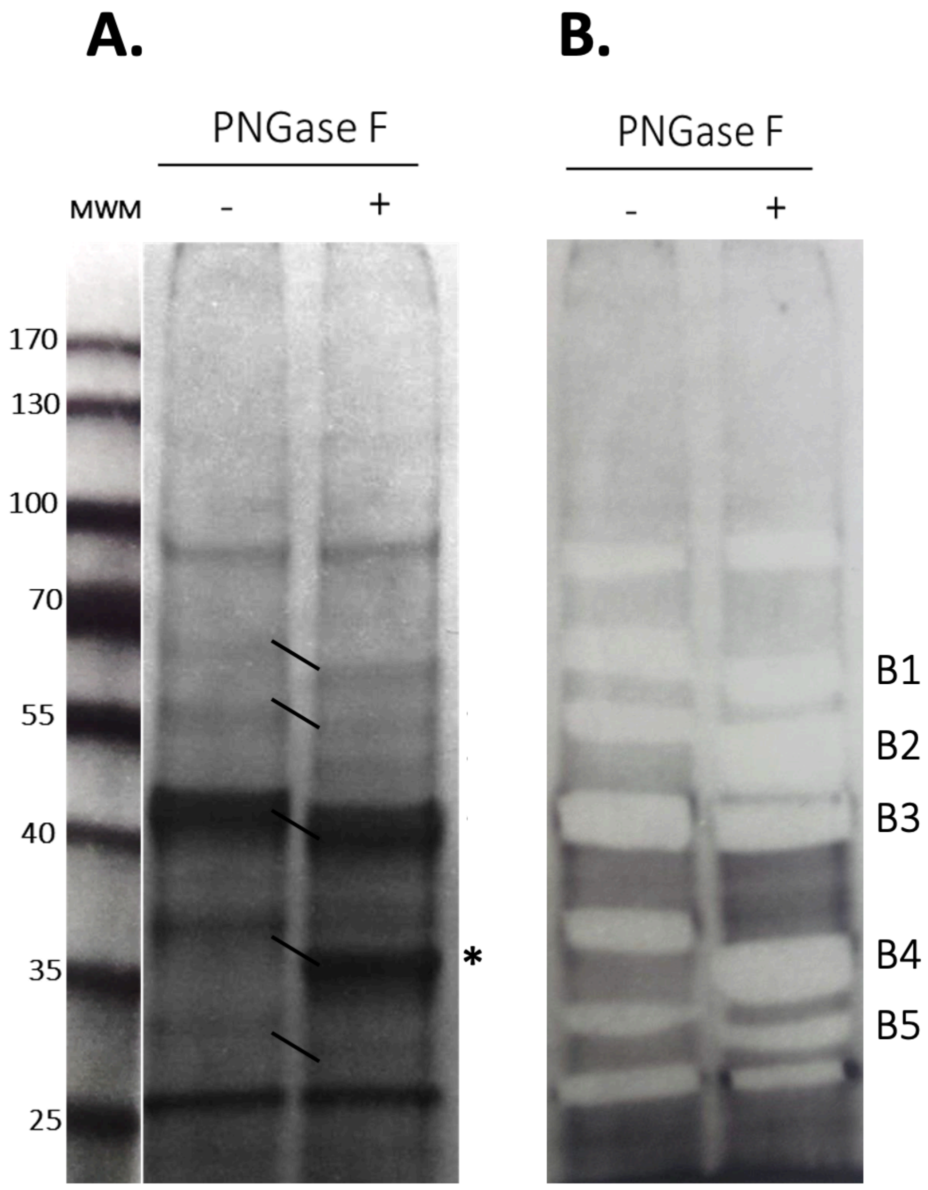


**Supplementary Figure S3. Enzymatic cleavage of *Lu. longipalpis* salivary glycoproteins with PNGase F.** 15 µg of salivary proteins were incubated overnight with (+) and without (-) PNGase F to cleave *N*-glycans. Samples were resolved on a 12 % SDS-PAGE gel. A) The change in the migration of the bands after loss of the *N*-glycans is visible. B) Bands susceptible to cleavage were excised and proteins identified by LC-MS. Excised bands with potential glycoproteins have been labelled B1-B5 (correspond to Supplementary Table 2). Asterisk indicates PNGase F enzyme band.

**A.**

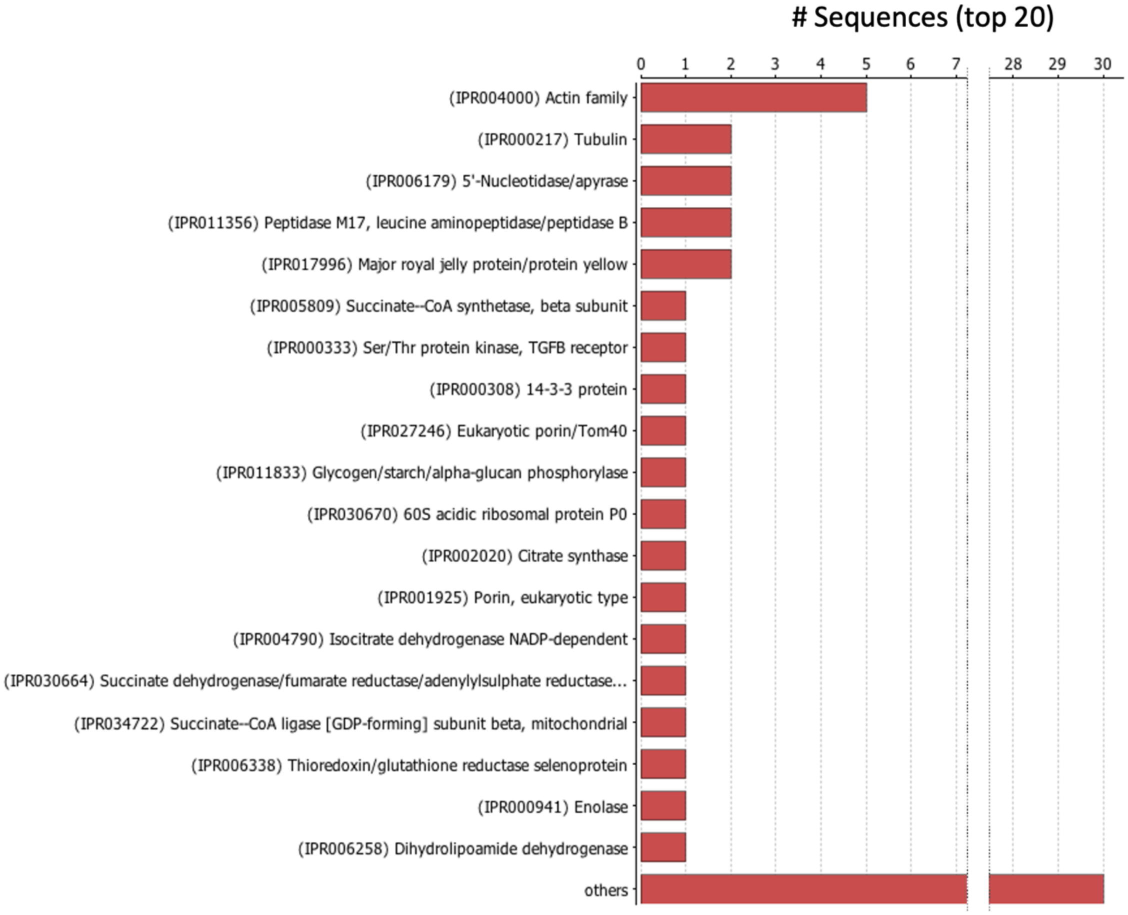

### Sequences (top 20)
0
1
2
3
4
5
6
7
28
29
30
(IPR004000) Actin family
(IPR000217) Tubulin
(IPR006179) 5'-Nucleotidase/apyrase
(IPR011356) Peptidase M17, leucine aminopeptidase/peptidase B
(IPR017996) Major royal jelly protein/protein yellow
(IPR005809) Succinate--CoA synthetase, beta subunit
(IPR000333) Ser/Thr protein kinase, TGFB receptor
(IPR000308) 14-3-3 protein
(IPR027246) Eukaryotic porin/Tom40
(IPR011833) Glycogen/starch/alpha-glucan phosphorylase
(IPR030670) 60S acidic ribosomal protein P0
(IPR002020) Citrate synthase
(IPR001925) Porin, eukaryotic type
(IPR004790) Isocitrate dehydrogenase NADP-dependent
(IPR030664) Succinate dehydrogenase/fumarate reductase/adenylylsulphate reductase...
(IPR034722) Succinate--CoA ligase [GDP-forming] subunit beta, mitochondrial
(IPR006338) Thioredoxin/glutathione reductase selenoprotein
(IPR000941) Enolase
(IPR006258) Dihydrolipoamide dehydrogenase
others

**B.**

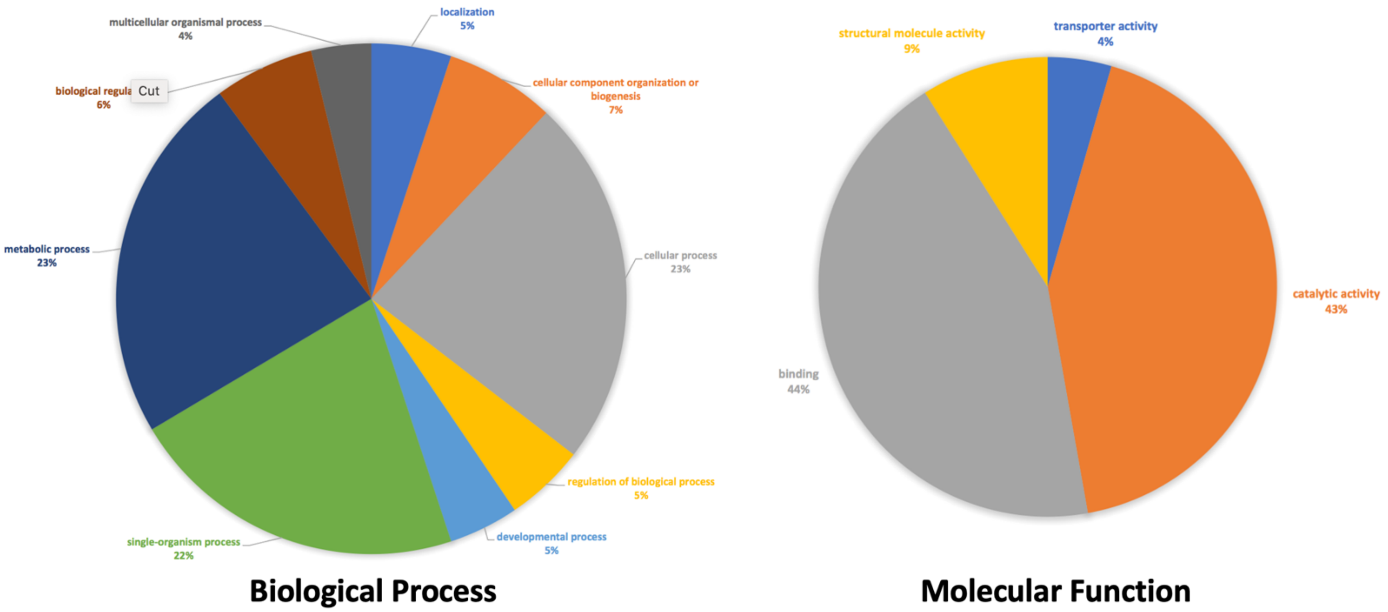

multicellular organismal process
4%
localization
5%
biological regula
6%
cellular component organization or biogenesis
7%
metabolic process
23%
cellular process
23%
regulation of biological process
5%
single-organism process
22%
developmental process
5%
Biological Process
structural molecule activity
9%
transporter activity
4%
catalytic activity
43%
binding
44%
Molecular Function


**C.**

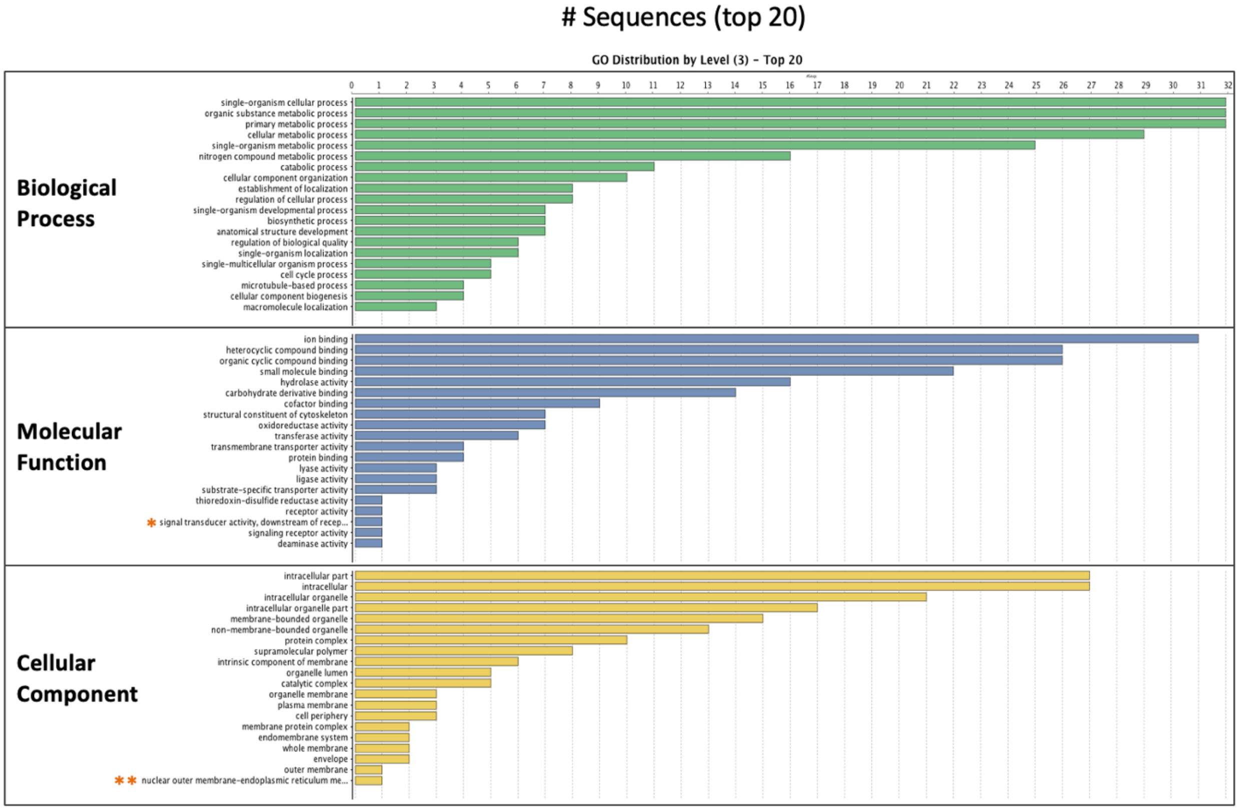

### Sequences (top 20)
GO Distribution by Level (3) - Top 20
Biological Process
single-organism cellular process
organic substance metabolic process
primary metabolic process
cellular metabolic process
single-organism metabolic process
nitrogen compound metabolic process
catabolic process
cellular component organization
establishment of localization
regulation of cellular process
single-organism developmental process
biosynthetic process
anatomical structure development
regulation of biological quality
single-organism localization
single-multicellular organism process
cell cycle process
microtubule-based process
cellular component biogenesis
macromolecule localization
Molecular Function
ion binding
heterocyclic compound binding
organic cyclic compound binding
small molecule binding
hydrolase activity
carbohydrate derivative binding
cofactor binding
structural constituent of cytoskeleton
oxidoreductase activity
transferase activity
transmembrane transporter activity
protein binding
lyase activity
ligase activity
substrate-specific transporter activity
thioredoxin-disulfide reductase activity
receptor activity
* signal transducer activity, downstream of recep...
signaling receptor activity
deaminase activity
Cellular Component
intracellular part
intracellular
intracellular organelle
intracellular organelle part
membrane-bounded organelle
non-membrane-bounded organelle
protein complex
supramolecular polymer
intrinsic component of membrane
organelle lumen
catalytic complex
organelle membrane
plasma membrane
cell periphery
membrane protein complex
endomembrane system
whole membrane
envelope
outer membrane
** nuclear outer membrane-endoplasmic reticulum me...
0 1 2 3 4 5 6 7 8 9 10 11 12 13 14 15 16 17 18 19 20 21 22 23 24 25 26 27 28 29 30 31 32

Supplementary Figure S4. **Analysis of candidate glycoproteins. A.** InterProScan protein family distributions. **B.** Classification of all glycoprotein candidates according to biological process and molecular function using Blast2GO. **C.** Classification of top 20 glycoprotein sequences according to biological process and molecular function using Blast2GO. *signal transducer activity, downstream of receptor, ** nuclear outer membrane-endoplasmic reticulum membrane network. The biological process classification revealed 86% of proteins are involved in various metabolic processes, including 30.23% in nucleobase-containing compound metabolic processes (GO:0006139), 23.26% in carbohydrate derivative processes (GO:1901135), and 23.26% purine-containing compound metabolic processes (GO:0072521). According to the molecular function classification, 81.25% of these proteins are involved in binding activities: 64.58% in ion binding (GO:0043167), 54.17% in heterocyclic (GO:1901363) and organic cyclic (GO:0097159) compound binding, and 45.83% in small molecule binding (GO:0036094). Additionally, 79.17% the glycoproteins were involved in catalytic activities, such as the 33.33% with hydrolase activity (GO:0016787). The cellular component classification shows 87.1% of the glycoproteins are intracellular (GO:0044424), of which 77.42% are cytoplasmic (GO:0005737) and 67.74% are located in intracellular organelles (GO:00043229).

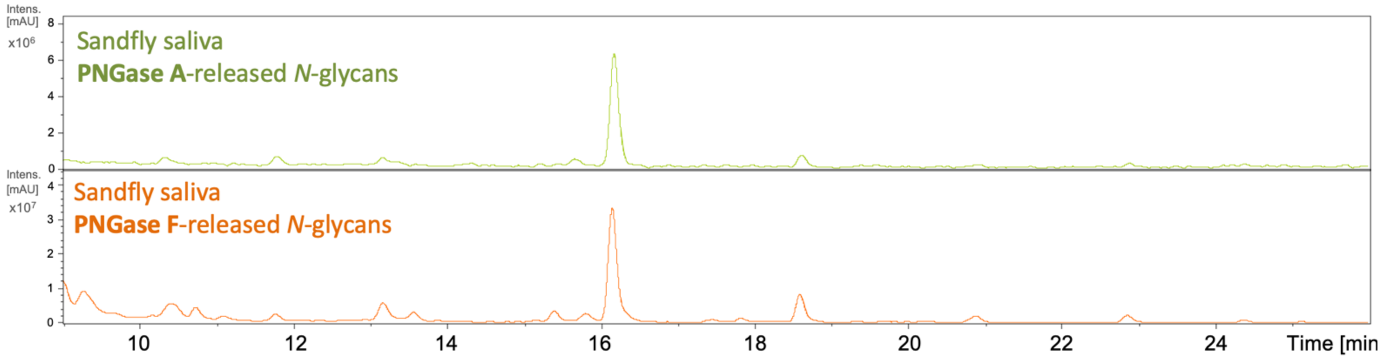


**Supplementary Figure S5.** HILIC-LC separation of PNGase A (top) and PNGase F (bottom) released *N*-glycans. Comparison after digestion with both enzymes indicates de absence of any α1,3 fucose modifications on the innermost GlcNAc of the trymannosyl core.

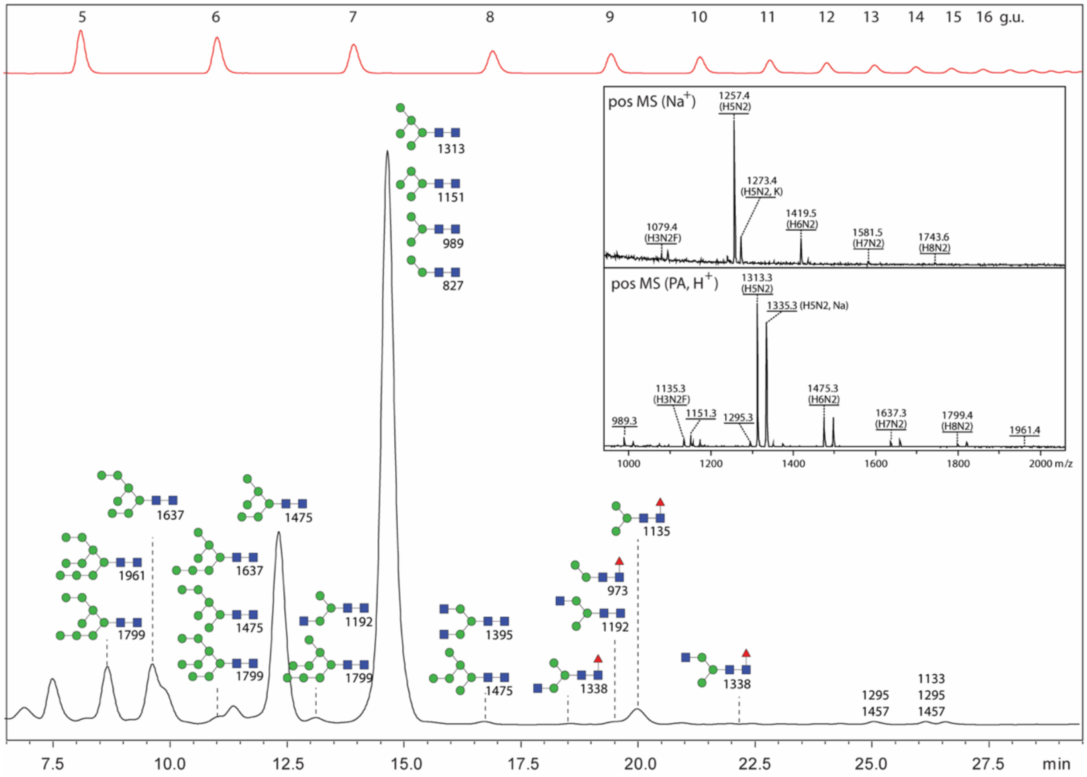


**Supplementary Figure S6. Analysis of pyridylaminated *N*-glycans from the sandfly by RP-amide HPLC and MALDI-TOF-MS.** PNGase F-released *N*-glycans were first subject to solid-phase extraction and analysed by MALDI-TOF-MS in positive mode (see inset, top panel; glycans detected primarily as $[M+Na]^+$ ions), prior to fluorescent labelling with 2-aminopyridine and analysis by MALDI-TOF-MS in positive mode (see inset, bottom panel; increase of 56 Da as compared to free glycans for the $[M+H]^+$ ions) and HPLC using an RP-amide column (main panel), pre-calibrated in terms of glucose units (GU). Each HPLC fraction was analysed by MS and MS/MS and, based also on comparison to previous studies using the same column, assignment of pauci- and oligomannosidic isomers was possible (annotations are with positive mode *m/z* values and the Symbolic Nomenclature for Glycans). Two late-eluting fractions (13 and 14 GU) contained putative *N*-glycans (*m/z* 1133, 1295 and 1457), but these could not be assigned by the comparative approach. Green circle, mannose; Blue square, N-Acetylglucosamine; Red triangle, fucose; Proc, procainamide.

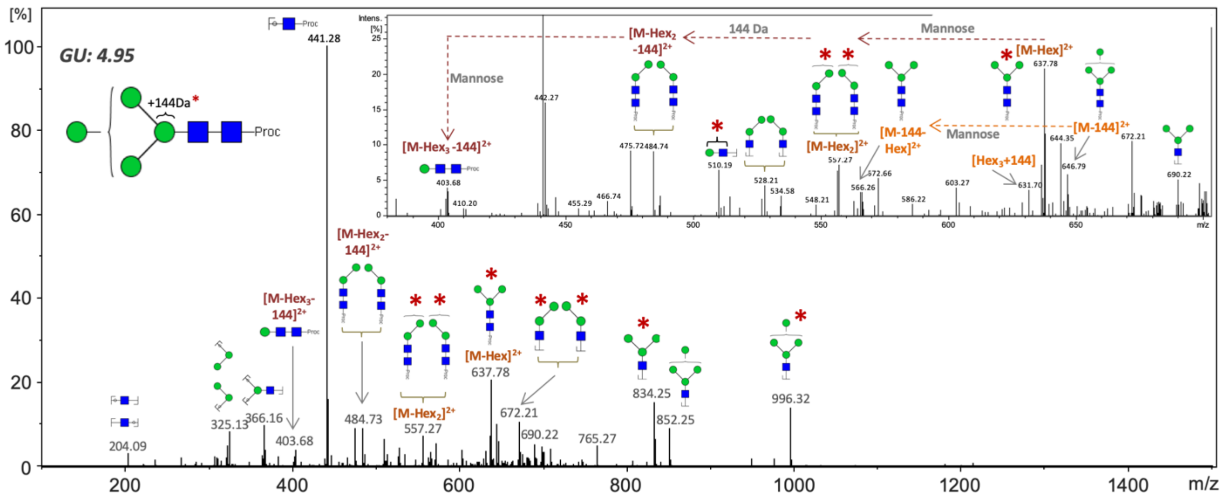


**Supplementary Figure S7. Positive-ion MS/MS fragmentation spectrum for glycan with *m/z* $[718.89]^{2+}$ corresponding a structure with a $Hex_4HexNAc_2$–Proc composition carrying a 144 Da modification.** The fragments with *m/z* 510.19 and 631.70 indicate the modification occurs on the inner most mannose of the chitobiose core and not the GlcNAc moiety. Asterisk denotes presence of the 144 Da modification on fragments. Green circle, mannose; Blue square, N-Acetylglucosamine; Proc, procainamide; GU, glucose units.

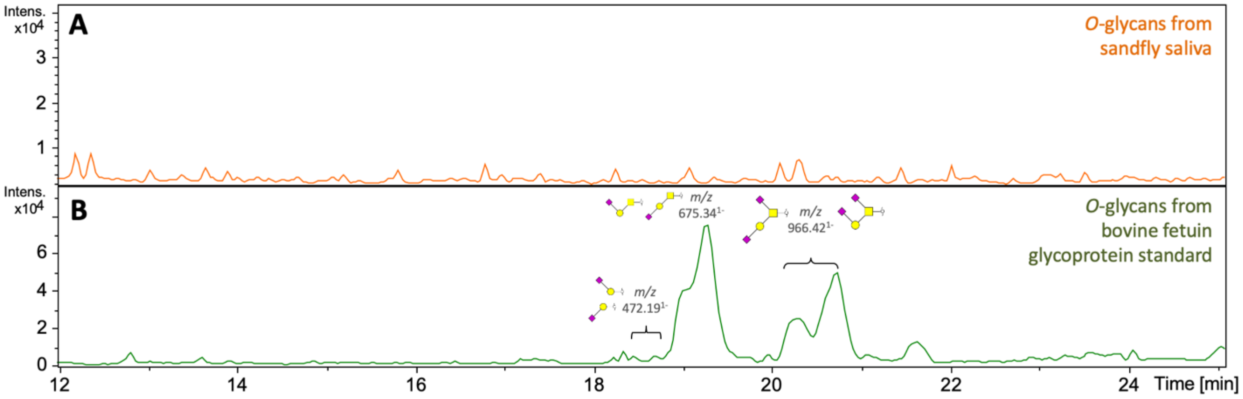


**Supplementary Figure S8. Beta-elimination release of *O*-glycans.** Released and reduced *O*-glycans were separated using a porous graphitised carbon column coupled with ESI-MS. No *O*-glycans were found in sand fly saliva (A); fetuin was used as a positive control (B). Yellow circle, galactose; yellow square, N-Acetylgalactosamine; Purple diamond, sialic acid.

A

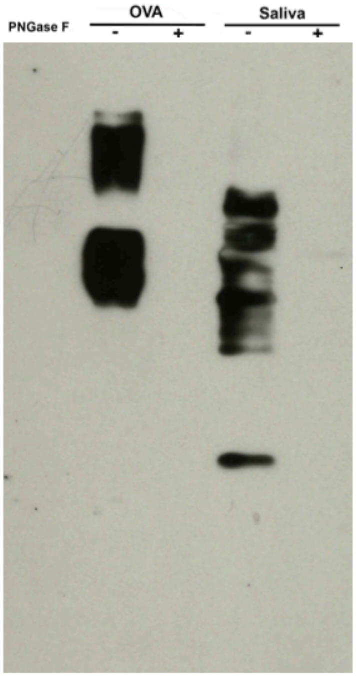


B

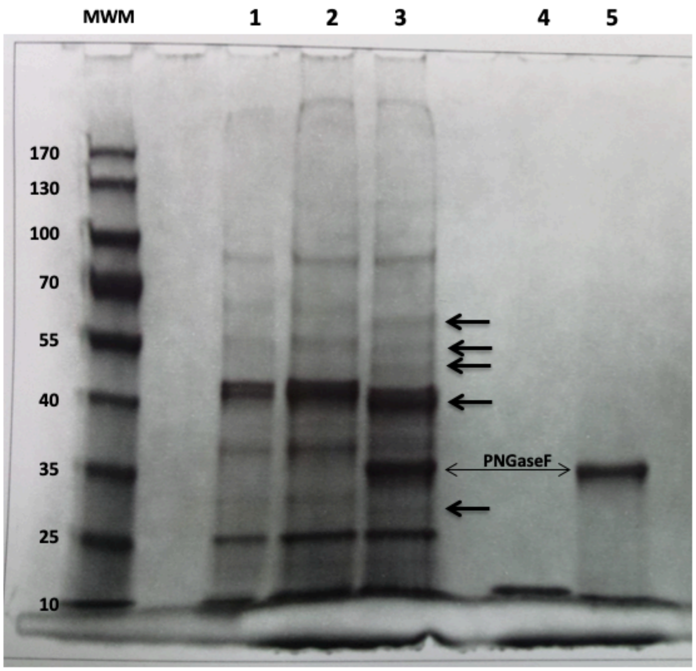


Supplementary Figure. (A) Full-length blot (Fig. 2) (B) Full-length Coomassie blue stained SDS-PAGE (Fig. S1)
